# Beyond Pulsing Dyes: Are Flickers the Language of the Mitochondrial Network?

**DOI:** 10.64898/2026.03.24.713912

**Authors:** Carolina Cierco, Francisco Santos, Sandrina Nóbrega-Pereira, Odete A.B. da Cruz e Silva, Diogo Trigo

## Abstract

Mitochondrial membrane potential (ΔΨm) is central to ATP production, ion homeostasis, and cell survival, reflecting the functional state of the inner mitochondrial membrane and oxidative phosphorylation. Accurate assessment of ΔΨm is therefore essential for understanding mitochondrial physiology and dysfunction in health, ageing, and disease. Lipophilic cationic fluorescent dyes, such as TMRM and TMRE, are widely used to monitor ΔΨm in live cells, enabling high-temporal-resolution imaging of both steady-state membrane potential and dynamic fluctuations. Beyond stable bioenergetic measurements, live-cell imaging reveals transient, reversible depolarisation events, known as mitochondrial “flickers.” These events, observed across multiple cell types and imaging platforms, are often associated with brief openings of the mitochondrial permeability transition pore (mPTP) and may represent regulated mitochondrial excitability, rather than irreversible damage. While excessive or synchronised depolarisations may signal mitochondrial injury, transient flickers are increasingly viewed as potential signalling mechanisms within the mitochondrial network. This work discusses methodological considerations for ΔΨm imaging, the biological significance of mitochondrial flickers, and the importance of distinguishing physiological events from probe- and light-induced artefacts, highlighting the emerging concept of mitochondria as dynamic and communicative bioenergetic networks.

## Introduction

Mitochondria are organelles with vital roles in cellular and organismal homeostasis. The highly folded inner mitochondrial membrane (IMM) houses the electron transport chain (ETC) and ATP synthase [1]. In the process of oxidative phosphorylation, electron transfer through complexes I–IV pumps protons (H^+^) from the mitochondrial matrix, generating an electrochemical gradient (ΔΨm) that drives ATP synthesis via ATP synthase [2]. ΔΨm is regulated by ATP synthase activity and proton leak/uncoupling and can be abruptly dissipated by opening of the mitochondrial permeability transition pore (mPTP) at contact sites, which also perturbs matrix Ca^2+^, pH, and mitochondrial volume [3]. ΔΨm supports ATP production, secondary active transport, and mitochondrial Ca^2+^ uptake through the mitochondrial calcium uniporter complex, contributing to ion homeostasis [4], [5].

Membrane architecture, fusion and fission dynamics [6], ΔΨm, and organelle contacts link mitochondrial bioenergetics with cell fate. [1], [3] Cardiolipin, the signature IMM phospholipid, supports negative curvature, stabilises ETC complexes and crista architecture, and participates in signalling during stress [3]. Changes in mitochondrial morphology, ΔΨm, and membrane permeability contribute to pathologies such as neurodegeneration [7] or cardiovascular disease [8]; moreover, ageing tissues feature more depolarised mitochondria, consistent with increased mPTP activity and bioenergetic failure [3].

Being so, assessing ΔΨm in experimental models offers a clearer insight into disease mechanisms, physiological processes, and theranostic targets. Several methods have consequently been developed to investigate mitochondrial membrane potential (ΔΨm).

Lipophilic cationic dyes such as tetramethylrhodamine methyl ester (TMRM), tetramethylrhodamine ethyl ester (TMRE), Rhodamine 123, DiOC6(3), or JC-1 accumulate in mitochondria according to the Nernst equation, reporting ΔΨm [4], [5]. These dyes are highly accessible and technically straightforward ubiquitous tools within the scientific community that facilitate real-time, live-cell imaging assessments of mitochondrial health and dynamics. Proper use of these probes requires non-quenching concentrations, and in times requires confirmation of mitochondrial localisation (e.g. using other mitochondrial dyes), normalisation for mitochondrial mass (in flow cytometry), and appropriate controls (such as FCCP or oligomycin, that selectively manipulate the membrane potential) [3].

### Live ΔΨm assessment

Live-cell imaging of mitochondrial function is thus an essential approach for resolving the dynamic behaviour of mitochondrial membrane potential (ΔΨm) and its transient fluctuations at the level of single mitochondria. Among the available tools, lipophilic cationic dyes, such as TMRM or TMRE, are the most widely used indicators for monitoring ΔΨm in live *in vitro* preparations [3], [9], [10] or *in vivo* [11]. These dyes accumulate electrophoretically within mitochondria in proportion to the membrane potential, following Nernstian principles, enabling high-temporal-resolution visualisation of both steady-state ΔΨm and rapid, localised depolarisation events [12].

The most common application is based on a non-quench (redistribution) mode, in which dyes are applied at low nanomolar concentrations, typically in the range of approximately 10–40 nM [3], [9], [10]. Under these conditions, mitochondrial fluorescence reflects Nernstian accumulation, and membrane depolarisation leads to a decrease in mitochondrial fluorescence, as the dye redistributes out of the matrix. Cells are usually equilibrated with the dye for at least 30 minutes at physiological temperature, and the dye is maintained in the imaging medium to preserve steady-state distribution [9], [12]. This mode is preferred for quantitative or comparative analyses, as fluorescence changes proportionally reflect ΔΨm variations [3], [13].

Whatever the experimental protocol, accurate ΔΨm imaging nonetheless requires rigorous control of dye handling and imaging conditions. Glassware is normally preferred, as nanomolar concentrations of TMRM/TMRE adsorb to polypropylene [14]; illumination should be minimised to the lowest power yielding a resolvable mitochondrial signal, to avoid detector saturation and excessive exposure [13], [15]. Confocal and two-photon microscopy are preferred, as optical sectioning and reduced out-of-focus excitation limit photodamage. Two-photon excitation, particularly, substantially reduces phototoxic ROS generation [3], [13], [15].

### Mitochondria Flickers

When applied under controlled conditions, TMRM- and TMRE-based live imaging reveals not only stable ΔΨm distributions, but also transient depolarisation events, sometimes termed mitochondrial “flickers” [16]. These are brief, localised ΔΨm losses that appear as transient decreases in mitochondrial fluorescence, in non-quench mode.

Flickers display heterogeneous kinetics and magnitudes [17], and can vary between mitochondria within the same cell. These events can be observed in virtually all live *in vitro* preparations (fig. 1), from human-derived (fig. 1a, supplementary video 1) and mouse embryonic (MEF) fibroblasts (fig. 1b, supplementary video 2), to neuronal primary cultures (fig. 1c, supplementary video 3) and cell lines (fig. 1d-e,, supplementary videos 4 and 5), labelled with TMRM (fig.1a, fig. 1c-e) or TMRE (fig. 1b). These are not restricted to an imaging setup, as they can be observed using different models of laser scanning confocal (fig.1a-d) or spinning disk (fig. 1e) microscopes. Although described to occur at the level of individual mitochondria (fig. 1c), a large portion of the whole mitochondrial network might reversibly depolarise (fig. 1a-b, fig.1 d-e).

**Fig. 1.**
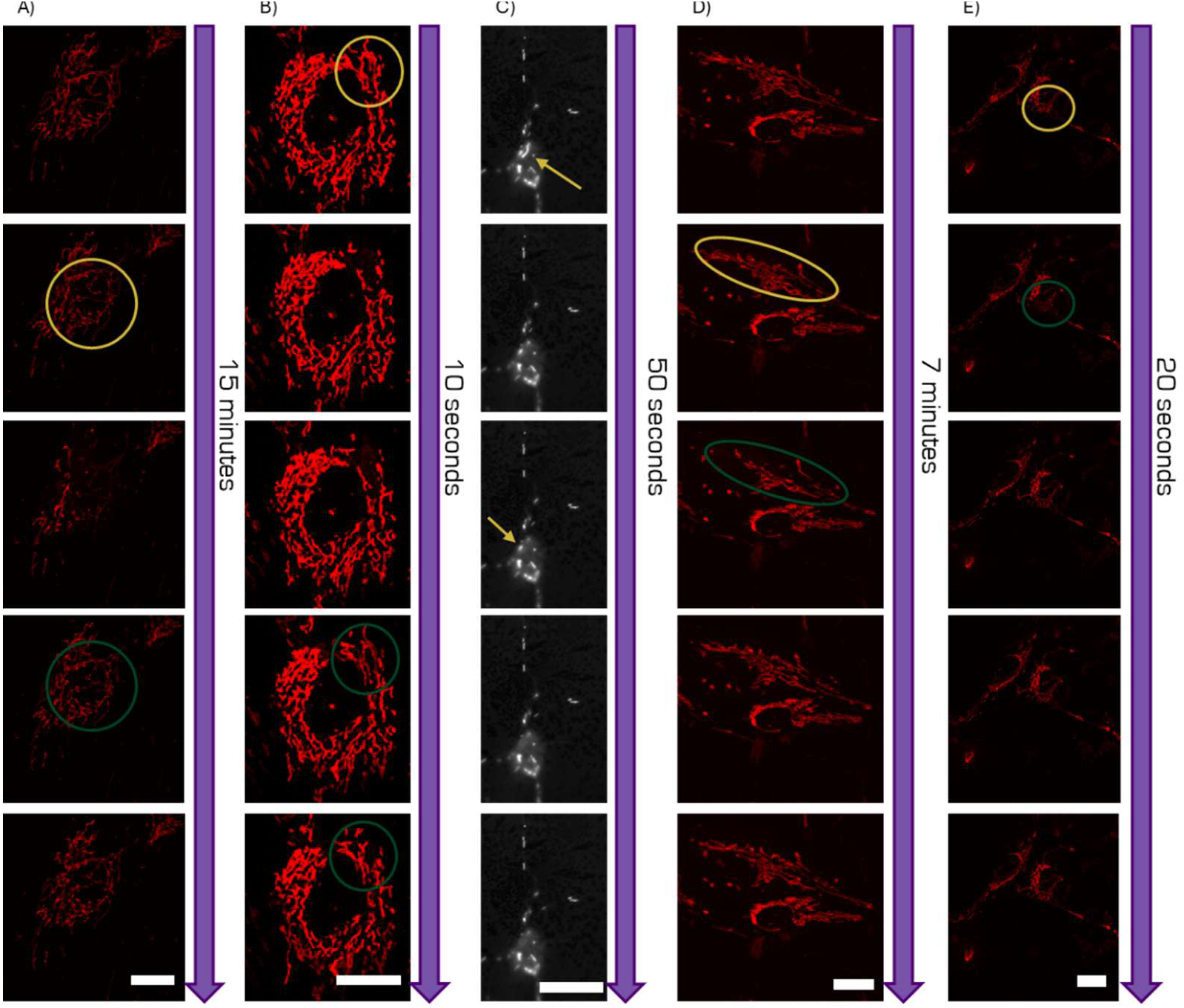
Examples of mitochondrial flicker in distinct preparations. A) TMRM stained human fibroblasts imaged by confocal microscopy. B) TMRE stained MEF imaged by confocal microscopy. C) TMRM stained cortical primary neuronal cultures imaged by confocal microscopy. D) TMRM stained SH-S5Y5 cells, imaged by confocal microscopy. E) TMRM stained SH-S5Y5 cells, imaged by spinning disk microscopy. Yellow indicates depolarising mitochondria, green indicates repolarising mitochondria. Scale bar is 20 µm.

These phenomena are less unequivocally identified in live animal imaging, due to the intricate three-dimensional architecture of the field and heterogeneous tissue environments, further complicated by respiratory- and cardiovascular-derived focal shifts derived focus (fig. 2, supplementary video 6).

**Fig. 2.**
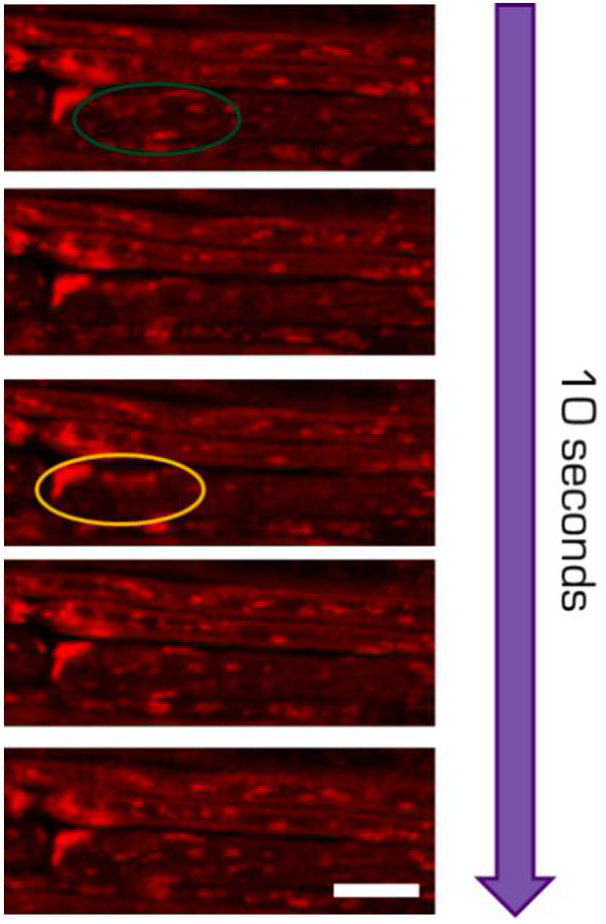
Possible mitochondrial flicker in mouse saphenous nerve. Saphenous fibres exposed in vivo in an anaesthetised mouse, labelled with TMRM and imaged by confocal microscopy. Yellow indicates depolarising mitochondria, green indicates repolarising mitochondria. Scale bar is 10 µm.

Although the challenges of investigating mPTP are well established [18], the flicker phenomenon is posited to be linked to transient, reversible openings of the mitochondrial permeability transition pore (mPTP), which allow momentary dissipation of the proton gradient without irreversible mitochondrial damage [19], [20]. Transient depolarisations may alternatively reflect H^+^ entry during ATP generation [21] or the localized exchange of contents between mitochondria [17], but the most common biological explanation seems to be transient mPTP flickering. These brief openings could also allow for the release of ions and small solutes, which may serve as a safety valve to prevent permanent damage or regulate matrix Ca^2+^.

A parallel, possibly similar phenomenon, termed superoxide flash, has been observed when researching mPTP activity [22]. Flash frequency and amplitude seem to be, at least partially, modulated by mitochondrial calcium uptake, local ROS levels, and the gating behaviour of the mPTP, particularly through cyclophilin D–dependent regulation, and can be artificially induced or amplified by imaging itself [19], [22], [23]. Photon-generated ROS arising from excessive illumination are hypothesised to trigger mPTP opening, creating depolarisation events reflecting experimental artifacts, rather than physiological signals; flash frequency seems to be reduced by antioxidants such as N-acetylcysteine and mPTP inhibition with cyclosporine A, under low-illumination conditions [20], [22].

Biologically, mitochondrial flickers and flashes are increasingly viewed as manifestations of an intermittent, regulated mitochondrial excitability process. Rather than representing pathological failure, transient mPTP openings and associated ΔΨm and redox signals may serve as localised signalling events that couple mitochondrial metabolism to intracellular communication pathways, including retrograde signalling to the endoplasmic reticulum and nucleus. The frequency of flickers and flashes correlates with metabolic demand, oxidative stress, and early mitochondrial dysfunction, and increases during conditions such as ischemia-reperfusion or metabolic overload. Sustained or synchronised events, however, may mark the transition from physiological signalling to mitochondrial injury and cell death [19], [23], [24].

Maintaining and regulating cellular homeostasis in response so internal and external stimuli requires a dynamic and adaptative mitochondria network [25]. Mitochondrial flickering represents a reversible depolarisation event that could act as a dynamic regulatory valve. These transients might be facilitating the stoichiometric exchange of ions, metabolites, or signaling factors, calibrating mitochondrial output to meet localized subcellular energetic demands.

## Conclusion

Live assessment of mitochondrial membrane potential using Nernstian cationic fluorescent dyes provides a powerful framework for dissecting mitochondrial homeostasis, while also demanding rigorous experimental discipline to distinguish genuine physiology from probe- and light-induced artefacts. These approaches reveal transient ΔΨm flickers and flashes, seemingly reflecting reversible mitochondrial permeability events, representing a biologically significant, albeit not entirely understood, layer of mitochondrial signalling. Beyond stochastic fluctuations, the wave-like flicker phenomena suggest an organised system of network-wide communication, where electrochemical signals could play a part coordinating vector for mitochondrial homeostasis within the complex cellular architecture.

## Material and methods

Commercial human (from the skin of the abdomen of a 25-year-old female donor) [26] or mouse embryonic fibroblasts [27] were cultured with Dulbecco’s Modified Eagle’s Medium supplemented with foetal bovine serum (FBS). Mouse primary cortical neurons, prepared as previously described [28], were cultured in neurobasal medium containing 2% B27 serum-free supplement, 2 mM l-glutamine, 1.5% glucose, penicillin (100 U/ml), and streptomycin (100 g/ml). SH-S5Y5 cells were cultured in equal parts of Minimal Essential Medium and F12 Ham’s F12 Nutrient Mix Medium, supplemented with 10% FBS. All preparations were plated in glass bottom dishes and incubated at 37°C in a humidified atmosphere of 5% CO_2_ and 95% air [29].

Imaging of human fibroblasts and SH-S5Y5 mitochondria was performed as previously described [29] by loading cells with 20 nM TMRM for 45 min prior to imaging using a Zeiss LSM 510 laser-scanning confocal microscope (Carl Zeiss, Oberkochen, Germany) with a × 63 oil-immersion Aprochromat objective. A similar SH-S5Y5 preparation was imaged using a Leica THUNDER Imager Cell Spinning Disk microscope with a × 63 oil-immersion objective. MEFs were incubated with 50 nM TMRE red for 20 minutes prior to washing and imaged using a Zeiss LSM 880 confocal microscope with a × 63 oil-immersion apochromat objective [27]. Primary neurons were loaded with 20 nM TMRM for 45 min, prior to imaging using a Zeiss LSM 700 laser-scanning confocal with a ×63 oil-immersion Aprochromat objective [28]

For *in vivo* imaging, the saphenous nerve was exposed as previously described [30] and labelled with TMRM (1 µM solution in sterile saline; pH=7), by soaking cotton balls with the TMRM solution and placing them over the entire exposed part of the nerve for 45 min [11]; the mouse was then placed on a modified microscope stage and imaged, using a confocal laser-scanning microscope Zeiss LSM 5 Pascal with a ×63 oil-immersion warmed (37°C) objective [30].

## Acknowledgements

We thank Leica Microsystems for providing the THUNDER Imager Cell Spinning Disk. The authors are grateful to Professor Kenneth J. Smith (University College London) for providing the raw data presented in Figure 2; these data were originally acquired by DT during doctoral research conducted under his supervision. Part of image acquisition was performed in the LiM facility of iBiMED, a node of PPBI (Portuguese Platform of BioImaging): POCI-01-0145-FEDER-022122. This work was funded by COMPETE2030-FEDER-00755200 and CI21-00276, and supported by Instituto de Biomedicina (iBiMED) - (UID/04501/2025, https://doi.org/10.54499/UID/04501/2025). DT is funded by CEECIND/CP1589/CT0025.

## Declaration of generative AI and AI-assisted technologies in the manuscript preparation process

During the preparation of this work the authors used Gemini and Grammarly to improve language, grammar, and readability. After using this tool/service, the authors reviewed and edited the content as needed and take full responsibility for the content of the published article.

## Notes

### Competing Interest Statement

The authors have declared no competing interest.

